# Subcellular proteomic profiling of human skeletal muscle reveals exercise-induced coordinated and compartment-specific protein remodeling

**DOI:** 10.64898/2026.02.08.703653

**Authors:** Elizabeth G. Reisman, Dale F. Taylor, Dingyi Yu, Cheng Huang, David J. Bishop, Nikeisha J. Caruana, John A. Hawley, Nolan J. Hoffman

**Affiliations:** Centre for Human Metabolism and Performance, Mary MacKillop Institute for Health Research, Australian Catholic University, Melbourne, Australia; Institute for Health and Sport, Victoria University, Melbourne, Australia; Mass Spectrometry Facility, St Vincent’s Institute of Medical Research, Fitzroy, Australia; RNA Mass Spectrometry Platform, Department of Biochemistry and Molecular Biology, Biomedicine Discovery Institute, Monash University, Clayton, Australia; Department of Biochemistry and Pharmacology, Bio21 Molecular Science and Biotechnology Institute, The University of Melbourne, Melbourne, Australia; Department of Sport and Exercise Sciences, Manchester Metropolitan University Institute of Sport, Manchester, United Kingdom

**Keywords:** Proteome, mitochondria, nucleus, cytosol, HIIT, endurance training

## Abstract

Exercise training induces extensive protein modifications in skeletal muscle, yet how acute exercise and training-induced molecular responses are spatially coordinated across muscle subcellular compartments remains unclear. Using subcellular fractionation combined with data-independent acquisition mass spectrometry, we profiled skeletal muscle mitochondrial, nuclear, and cytosolic proteomes in response to an acute bout of intense cycling (pre-, mid-, post- and 3 h post-exercise) and after eight weeks of endurance training in 40 healthy adults (20 males and 20 females). Acute exercise triggered coordinated, compartment-specific proteomic remodelling, including reductions in protein translation and import machinery concomitant with increased redox-related proteins. Notably, acute exercise increased markers of ribosomal translation within the mitochondrial fraction, revealing ribosomal scaffold protein RACK1 as a potential regulator of subcellular translational control under contractile stress (confirmed by targeted immunoblotting). The nuclear proteome displayed transient remodelling of RNA-processing and chromatin-associated proteins, while cytosolic changes were modest. Endurance training induced robust proteomic remodelling across all compartments, including increased markers of mitochondrial oxidative metabolism and proteostasis. While there were sex differences at baseline, subcellular proteomic responses were largely conserved between sexes. We provide the first comprehensive, time-course subcellular characterisation of the skeletal muscle proteome, revealing regulation of translational machinery underlying the acute exercise response.

## Introduction

Exercise is a potent lifestyle intervention for improving metabolic, cardiovascular, neurological, and overall health, reducing the risk of numerous chronic diseases ^1, 2, 3^. Although its systemic health and clinical benefits are well established, the molecular mechanisms through which exercise remodels human tissues, particularly skeletal muscle, are not fully understood. Skeletal muscle is a dynamically regulated and metabolically active tissue, playing a central role in locomotion, glucose homeostasis and whole-body energy balance ^4, 5^. Exercise represents a robust physiological stimulus that disrupts whole-body and cellular energy homeostasis, with contracting skeletal muscles triggering widespread molecular responses across multiple cells and tissues in response to the increased metabolic and oxygen demands ^6^. However, how acute exercise-induced perturbations reorganise protein networks in tissues such as skeletal muscle, and how these responses accumulate to drive long-term adaptation in response to endurance training, remain incompletely defined.

Despite advances in transcriptomic and whole-tissue proteomic profiling ^7, 8, 9, 10^, relatively little is known about how exercise remodels protein networks within distinct subcellular compartments of human skeletal muscle, and proteomic spatial resolution evidence remains limited ^11, 12, 13, 14^. Protein functionality and regulation is highly dependent on subcellular localisation with compartment-specific proteostasis including protein synthesis, folding, degradation, and import central to maintaining muscle adaptation and function under stress ^15^. Organelles such as the mitochondria, nuclei, and cytosol engage distinct, localised regulatory programs, yet interact dynamically through protein trafficking, metabolite signalling, and stress-responsive proteome remodelling mechanisms ^16, 17, 18, 19, 20, 21^. Notably, contractile-induced transcriptomic responses often fail to predict corresponding protein changes ^22, 23^, underscoring the need for direct measurement and subcellular spatial resolution of protein abundance. Prior studies have used univariate approaches or analysis of only a single subcellular fraction, limiting insights into potential coordinated, intracellular protein-level changes occurring across multiple subcellular compartments ^24, 25, 26^. Whole-tissue proteomic analyses are further complicated due to the vast abundance of sarcomeric proteins, which obscures detection of lower-abundance proteins residing in compartments such as the mitochondria and nuclei with potential regulatory and functional significance ^7, 27^. As a result, the subcellular molecular machinery underlying acute exercise and exercise training responses, and their coordination across organelles remains largely unexplored.

Emerging evidence suggests that subcellular protein regulation and inter-organelle communication are critical for exercise training-induced adaptation, including mechanisms of mitochondrial–nuclear signalling, ribosomal localisation, localised intracellular stress responses, and coordinated regulation of protein quality control pathways ^20, 28, 29, 30, 31^. However, there have been no comprehensive proteome-wide, time-course investigations of how subcellular protein abundance changes in response to exercise in human muscle. The potential contribution of biological sex to these molecular processes also remains underexplored, and while sex differences in exercise metabolism and substrate utilisation are recognised, it is unclear whether such differences result in exercise-induced compartmental proteomic regulation. To address these knowledge gaps, we performed the first subcellular, global proteomic analysis of fractionated human skeletal muscle from both males and females, quantifying exercise-induced changes in protein abundance across mitochondrial, nuclear, and cytosolic compartments, enabling characterisation of temporal and subcellular remodelling of protein in response to both acute exercise and following eight weeks of exercise training (**Figure 1a**).

**Figure 1 -.**
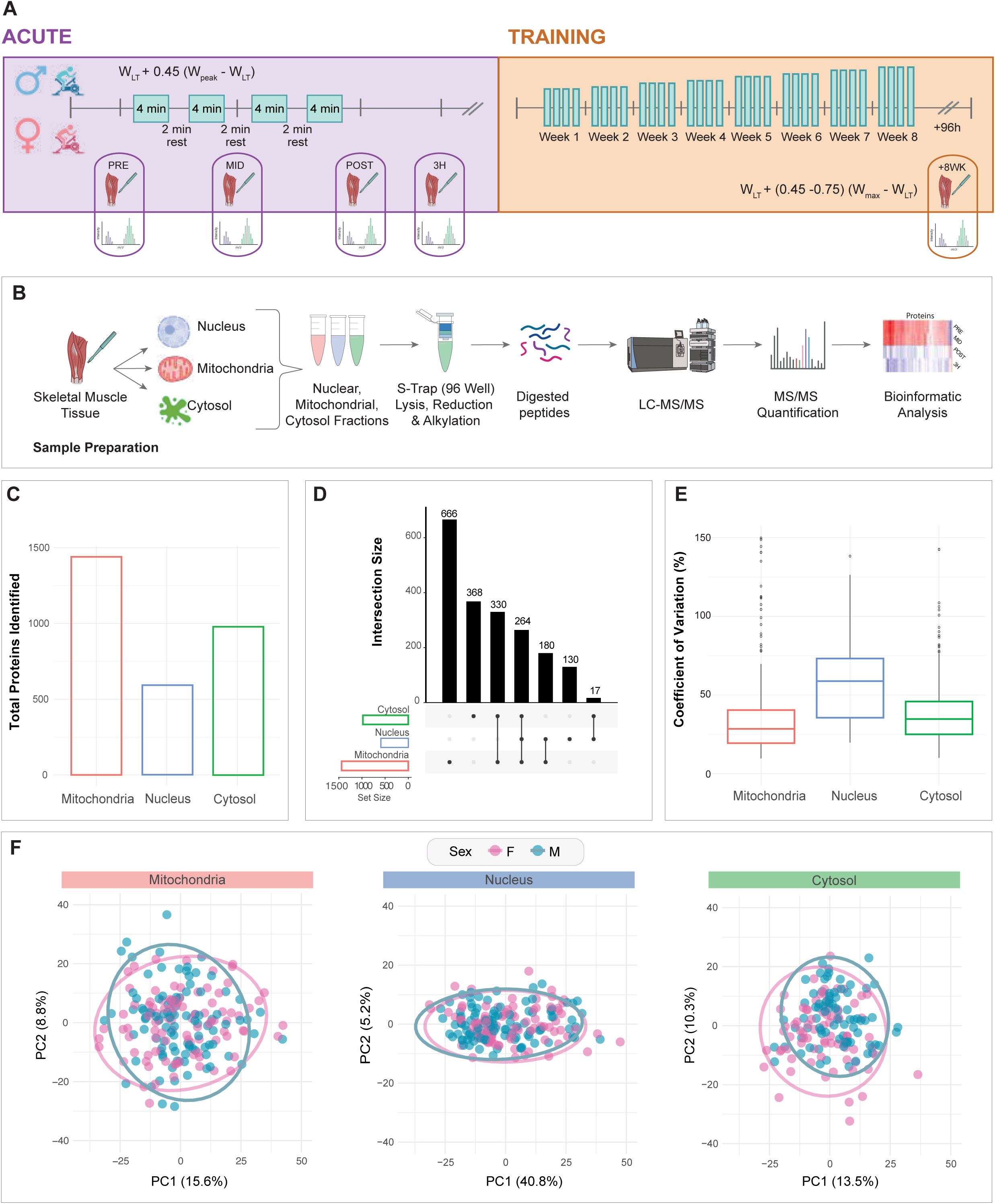
Overview of acute exercise and training protocols and subcellular fractionation and MS analysis workflows. **a)** Overview of the overall study protocol for males and females including experimental exercise session and 8-week exercise training period (with the post-training biopsy collected 96 h after the last exercise bout). The first High-Intensity Interval Training (HIIT) session and associated timepoints (PRE, MID, POST & +3 h) of muscle biopsy collection are specified for mass spectrometry (MS)-based subcellular proteomics analysis. Each bar represents a single exercise session. Created with BioRender.com **b)** Human muscle biopsy sample preparation and analysis workflow using mass spectrometry-based proteomics: Skeletal muscle samples were subjected to subcellular fractionation and processed for proteomic analysis. Samples were digested into peptides, which were analysed by liquid chromatography–mass spectrometry (LC-MS/MS). Peptide data were acquired using data-independent acquisition (DIA) mode for protein identification and quantification and analysed using a suite of bioinformatic tools. Created with BioRender.com **c)** Total number of proteins identified and quantified in all three subcellular fractions following data clean-up filtering and normalisation. **d)** UpSet plots summarise proteins identified/quantified in each subcellular fraction and in multiple fractions. The bottom left horizontal bar graph indicates the total number of proteins per subcellular fraction. The circles in each panel’s matrix represent the unique proteins within a single subcellular fraction, and connected circles indicate overlapping proteins between the subcellular fractions. The top bar graph in each panel summarises the number of unique or overlapping proteins for each comparison. **e)** Coefficient of variations within each fraction representing the variability of all proteins identified and quantified between samples. Each box represents the interquartile range (IQR) and median of the percentage of the coefficient of variation. Each whisker represents 1.5 * IQR for both directions. Outliers are displayed. **f)** Principal component analysis of sex for all samples in each subcellular fraction, with each dot representing a unique biological sample (female samples shown in pink; male shown in blue).

## Results

### Human muscle subcellular proteomic dataset overview

Subcellular proteomic data are reported from the mitochondrial, nuclear, and cytosolic fractions of *vastus lateralis* muscle biopsies collected from 40 healthy, untrained adults (20 males, 20 females) across an acute bout of high-intensity interval training (HIIT; i.e., before (PRE), during (MID), immediately after (POST), and 3 h-post (3H)) and following an 8-week HIIT intervention (see Table S1, Tab 1 for participant characteristics). Subcellular fractionation was employed to overcome the large dynamic range of skeletal muscle protein abundance and enhance detection of lower-abundance proteins within each fraction. We combined an optimised fractionation workflow with liquid chromatography-mass spectrometry (LC–MS/MS) analysis and data independent acquisition (DIA) (**Figure 1b**), enabling high-throughput profiling of this complex proteomic dataset from a large human muscle sample set. Across all samples, we detected 2,199 total proteins in the mitochondrial fraction, 1,249 in the nuclear fraction, and 1,882 in the cytosolic fraction (1% FDR; Table S1, Tabs 2–4). After removing proteins with >30% missing values in each fraction, the filtered dataset comprised 1,440 mitochondrial, 591 nuclear, and 979 cytosolic fraction proteins for downstream analysis (**Figure 1c**; Table S1, Tabs 5–7). Of the 1,440 total proteins detected in the mitochondrial fraction, 49% were mitochondrial-annotated according to the curated MitoCarta3.0 database ^32^, suggesting detection of potential contaminant proteins from subcellular fractionation and/or proteins that localise in multiple subcellular compartments or dynamically associate with mitochondria in skeletal muscle in response to exercise.

Relative purity across subcellular fractions was confirmed by assessing known resident proteins from each fraction (**Supp. Figure 1a–b**; Table S1, Tab 8). Immunoblot analysis of these protein markers demonstrated mean fraction purities of 81.8% (mtHSP70; mitochondrial), 80.4% (H3; nuclear), and 84.9% (LDHA; cytosolic). The mitochondrial fraction contained the greatest number of unique proteins (666), and shared the largest overlap with the cytosolic fraction (330) (**Figure 1d)**, consistent with known bidirectional protein translocation and shared cellular metabolic regulatory functions between these compartments at rest, during and/or following exercise ^33^.

Initial quality control of the processed, filtered protein abundance matrix was assessed by calculating protein-level coefficients of variation (CVs) within each fraction (**Figure 1e**). Median CVs of 29.2 (mitochondria), 34.8 (cytosol), and 58.9 (nucleus) fell within expected ranges for label-free quantitative proteomics ^34, 35^, with mitochondrial and nuclear protein abundance respectively exhibiting the lowest and highest variability across the three fractions. These muscle subcellular data indicate sufficient fractional purity and quality of proteomic data to enable reliable differential expression analyses. Principal component analysis (PCA) of the three subcellular proteomes showed variance distributed across multiple dimensions (i.e., PC1 and PC2) and no clear clustering by sex (**Figure 1f**).

### Acute Exercise-Induced Changes in Subcellular Protein Abundance

To assess compartment-specific proteomic responses to an acute bout of exercise, PCA was performed on mitochondrial, nuclear, and cytosolic fractions generated by LC–MS/MS (**Figure 2a**) revealing distinct fractional and temporal changes in protein abundance (**Figure 2b**). The mitochondrial proteome showed the largest temporal regulation, particularly from PRE to POST (PC1 = 18%), reflective of rapid responses to acute exercise-induced energetic and oxidative stress. The nuclear fraction displayed a relatively delayed but sustained shift POST and 3H, consistent with transcriptional, RNA-processing, and protein-folding responses, whereas the cytosolic fraction exhibited more modest exercise-induced alterations confined largely to POST and 3H. Mean subcellular protein intensities (**Figure 2c**) further reflected these temporal patterns. Mitochondrial-annotated proteins from MitoCarta 3.0 ^32^ robustly decreased POST (P = 1.19 × 10□□) and partially recovered after 3H, whereas nuclear proteins annotated in the Human Protein Atlas (HPA) ^36^ decreased only at 3H (P = 2.5 × 10□□). Cytosolic HPA-annotated proteins remained relatively stable, consistent with more mitochondrial/nuclear relative to cytosolic protein remodelling across acute exercise timepoints.

**Figure 2 -.**
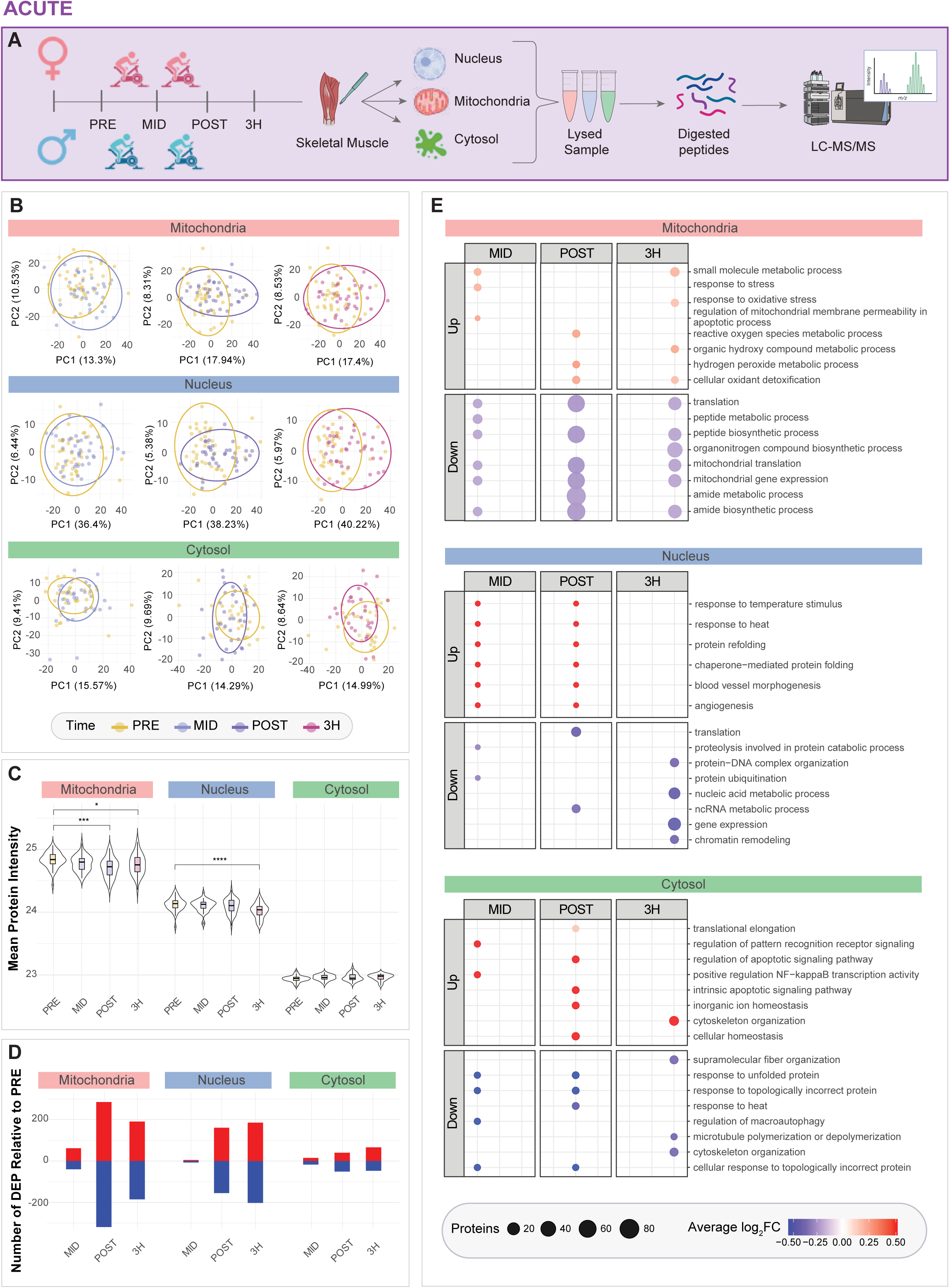
Changes in subcellular protein abundance in response to acute exercise. **a)** Overall workflow for analysing mitochondrial, nuclear and cytosolic fractions isolated from skeletal muscle of both males and females via LC-MS/MS. Created with BioRender.com **b)** Principal component analysis of samples per time point in the mitochondrial, nuclear and cytosolic fraction relative to respective pre-exercise baseline. **c)** Violin plots showing mean mitochondrial, nuclear and cytosolic protein content across acute timepoints (PRE, MID, POST and +3 H). Proteins were annotated as mitochondrial according to Mitocarta 3.0 ^32^ and nuclear or cytosolic according to the Human Protein Atlas^36^. Within each violin plot, the box shows the interquartile range (IQR) and median, and the whiskers extend to 1.5 × IQR. Outliers are not displayed. *P ≤ 0.01, ** P ≤ 0.001, *** P ≤ 0.0001 based on ANOVA with Tukey post-hoc testing for pairwise timepoint comparisons. **d)** Number of differentially expressed proteins of the mitochondrial, nuclear and cytosolic fractions according to *limma* analysis with Benjamini–Hochberg (BH) adjustment, using an adjusted P-value < 0.05, highlighting proteins that are either upregulated (red) or downregulated (blue) per acute exercise timepoint as determined by their log_2_ fold change. **e)** EnrichPlot dot plot analysis of differentially expressed proteins at each acute timepoint, annotated as mitochondrial (Mitocarta 3.0 ^32^) or as nuclear/cytosolic (Human Protein Atlas ^36^). Pathway enrichment was performed using Gene Ontology Biological Processes (GOBP). For each timepoint, the top eight enriched terms (ranked by adjusted P-value) are shown for upregulated and downregulated proteins. Dot size represents the number of proteins associated with each term, while colour saturation reflects the average log□ fold-change for that timepoint and direction of regulation. A full list of enriched terms is provided in Supplementary Table 2.

Differential expression analysis (**Figure 2d)** aligned with these overall proteomic signatures whereby mitochondrial proteins showed the greatest acute exercise remodelling, with extensive changes POST (285 up-regulated, 319 down-regulated) and 3H (191 up, 185 down), while MID exhibited the fewest changes (62 up, 40 down). These patterns are consistent with dynamic subcellular protein regulation at the mitochondria in response to acute exercise-induced energetic stress ^37, 38^. These differentially expressed proteins (DEP) in the mitochondrial fraction included transport and metabolic regulatory proteins such as SLC25A46, SLC2A4, and ATP5IF1 at MID; MRPL49, TIMM21, and the iron-binding protein LTF at POST; and stress-sensitive proteins including RYR1 and HSPA1A at 3H. Nuclear responses occurred mostly POST and 3H (POST: 155 up, 161 down; 3H: 186 up, 202 down), featuring coordinated regulation of splicing-related heterogeneous nuclear ribonucleoproteins (HNRNPs; HNRNPL, HNRNPU, HNRNPA2B1), ribosomal proteins (RPLP2, RPL13A, RPL26), and heat shock proteins (HSPB1, HSPB6, HSPB7, HSPB8). Notably, HNRNP family members are key regulators of alternative splicing, aligning with recent transcriptomic evidence that exercise induces widespread RNA-processing and splicing changes in human skeletal muscle ^39^. Cytosolic changes were smaller in magnitude (POST: 40 up, 51 down; 3H: 66 up, 47 down). Across all fractions, log□ fold changes were relatively modest in magnitude (typically within ± 0.5 versus PRE), suggesting broad but low-magnitude subcellular proteomic remodelling across compartments in response to acute exercise (Table S2) ^40, 41^.

Gene ontology biological process (GOBP) enrichment of compartment-annotated DEP highlighted distinct biological pathway regulation and compartmental responses to exercise (**Figure 2e**). For example, exercise led to upregulation of oxidative and metabolic stress responses in the mitochondrial fraction, including enrichment of pathways involved in oxidant detoxification (GO:0098869), hydrogen peroxide metabolic process (GO:0042743), and broader reactive oxygen species metabolic processes (GO:0072593). These changes occurred alongside robust downregulation of mitochondrial translation (GO:0032543) and mitochondrial gene expression (GO:0140053), suggesting a shift in mitochondrial biological process regulation from protein synthesis toward redox regulation and metabolic control during/post-acute exercise, consistent with the transient post-exercise reduction in translational activity reported in human skeletal muscle ^42, 43^. Nuclear pathway responses to acute exercise reflected stress-adaptive remodelling, characterised by downregulation of core gene regulatory processes, including translation (GO:0006412), gene expression (GO:0010467), nucleic acid metabolic processes (GO:0090304), and chromatin remodelling (GO:0006338). In contrast, in response to acute exercise, the cytosolic fraction demonstrated upregulation of translation elongation (GO:0006414), cytoskeletal organisation (GO:0007010), and apoptotic and inflammatory signalling (including NF-κB-related pathways), while pathways related to unfolded protein handling, microtubule dynamics, autophagy regulation, and heat stress were transiently downregulated in the cytosolic fraction (**Figure 2e**).

Together, these results demonstrate that a single bout of exercise induces rapid and compartment-specific proteome restructuring in human skeletal muscle, with mitochondrial fraction proteins responding earliest, nuclear fraction proteins adapting later, and the cytosolic fraction proteome remaining relatively stable. Despite these temporal differences, overall fold-changes were modest, likely reflecting tightly regulated subcellular protein abundance between compartments. Sex did not have a significant influence on DEP patterns within any fraction, despite subcellular proteome differences detected between sexes at baseline consistent with established sex differences in substrate utilisation (e.g., higher mitochondrial lipid-oxidation protein abundance in females and greater glycogen-metabolism/NAD□-related protein abundance in males; (**Supp. Figure. 2a -b**; Table S5).

### Eight weeks of HIIT induces skeletal muscle subcellular compartment-specific proteomic remodelling across mitochondrial, nuclear and cytosolic fractions

Endurance training is known to improve aerobic capacity and skeletal muscle mitochondrial content/function ^44, 45^, but how these changes manifest in terms of subcellular proteome abundance remains less well characterised. Following 8 weeks of HIIT, we profiled mitochondrial, nuclear, and cytosolic fractions using DIA-LC-MS/MS to examine chronic, compartment-specific protein remodelling (**Figure 3a**). Training resulted in the expected improvements in exercise capacity and whole-body physiology, including increased lactate threshold and maximal oxygen uptake (V□O□max) in both sexes. Mean improvements in lactate threshold (M = 26%, F = 32%) and V□O□max (M = 22 %, F = 16 %) are presented in **Figure 3b**.

**Figure 3.**
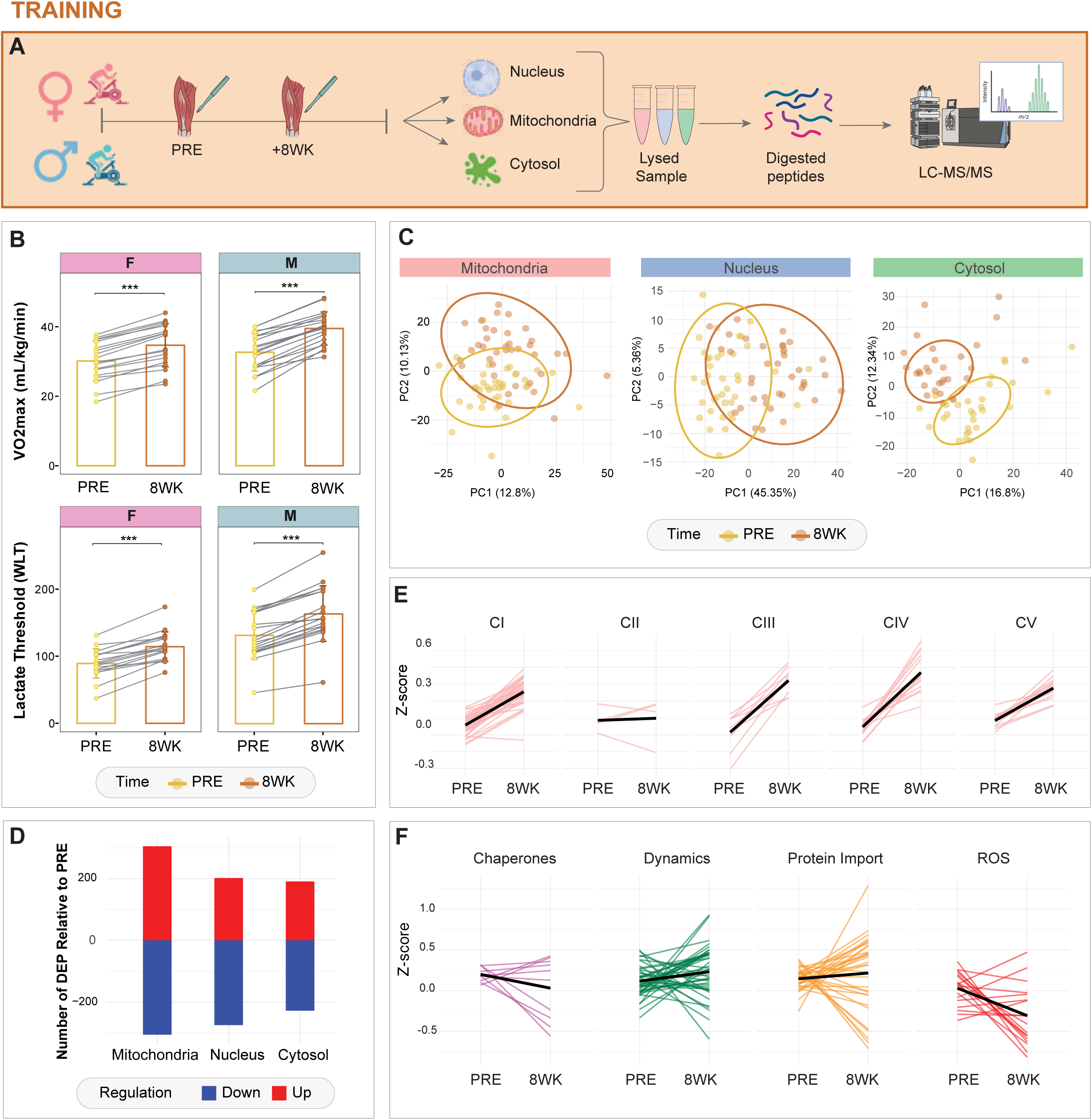
- Skeletal muscle subcellular proteomic adaptations to 8 weeks of endurance-based exercise training: proteomic profile changes across subcellular fractions. **a)** Overall workflow for isolation and LC-MS/MS analysis of subcellular fraction (mitochondrial, nuclear and cytosol) from skeletal muscle samples of both males and females collected before and following 8-weeks of endurance exercise training. **b)** Comparison of changes in whole-body physiological and blood lactate markers of aerobic capacity and exercise performance following 8 weeks of high-intensity interval training (HIIT) between males (M) and females (F). V□O2max = maximal oxygen consumption (WLT) = power at lactate threshold. *P ≤ 0.01, **P ≤ 0.001, ***P ≤ 0.0001 based on mixed two-way ANOVA with repeated measures to investigate the influence of sex and time. **c)** Principal component analysis of samples in each subcellular fraction following 8 weeks of training relative to pre- exercise. **d)** Number of differentially expressed proteins according to *limma* analysis and adjusted P values in each subcellular fraction that are either upregulated (red) or downregulated (blue) as determined by their log2 fold change. **e)** Scaled profile plots showing the relative abundance of mitochondrial protein pathways as annotated by MitoCarta 3.0 for each complex subunit (CI-CV) of the electron transport chain. with the mean relative abundance (black) of protein subunits in each complex group presented for PRE and post-8 weeks of HIIT (8WK). **f)** Mitochondrial pathways presented as scaled profile plots, including proteins associated with chaperone proteins (light purple), mitochondrial dynamics (dark green), protein import (orange) and reactive oxygen species (ROS) formation (red) as annotated by MitoCarta 3.0. Each pathway features a scaled profile plot showing the relative abundance of each mitochondrial protein annotated within the specific pathway for PRE and post 8 weeks exercise training, and the mean relative abundance (black) of the collective proteins in each group is also presented.

Based on the lack of sex differences observed in the subcellular proteome in response to exercise training (Table S5, Tab 1-3), PCA and differential expression analyses were performed across all 40 participants to further characterise compartment-specific adaptations in this larger cohort. PCA across all subcellular fractions (**Figure 3c**) showed clear separation of pre- versus post-training overall proteome profiles, with PC1 explaining 45% of the variance (PC2: 5%). Mitochondrial (PC1: 13%) and cytosolic (PC1: 17%) proteomes also exhibited shifts in their overall signatures before and after training, with PC1 contributing less to this variance (PC2: 10% and 12%, respectively) compared to the nuclear fraction. Differential expression analysis (**Figure 3d**) comparing protein abundance before and after 8 weeks of HIIT within each subcellular fraction revealed that the mitochondrial proteome underwent a balanced, robust response of upregulated (305) and downregulated (306) DEP (Table S3, Tab 2-4). In the mitochondrial fraction, HIIT induced upregulation of OXPHOS proteins, mitochondrial ribosomal biogenesis (e.g., large (MRPL) and short (MRPS) mitochondrial ribosomal proteins (mtRibo), and protein import machinery (e.g., TOMM and TIMM complexes). These changes are consistent with enhanced mitochondrial functional capacity and proteostasis following exercise training ^9, 46^. In the nucleus, 202 proteins were upregulated and 275 downregulated. The cytosolic fraction exhibited 191 upregulated and 228 downregulated proteins, reflecting widespread training-induced adaptations across all three compartments including fraction-specific protein responses. GOBP enrichment of upregulated proteins further supported these fraction-specific proteomic adaptations (**Supp. Figure 2c**), confirming increased mitochondrial oxidative phosphorylation, electron transport, and protein import pathways, enrichment of chromatin-associated and gene regulatory processes in the nuclear fraction, and enhanced biosynthetic and metabolic programs within the cytosol.

We next examined the relative abundance of subsets of proteins involved in specific mitochondrial pathways using scaled profile plots (**Figure 3e and 3f**), which revealed robust adaptations following exercise training. An upregulation of OXPHOS complexes CI–CV was observed within the mitochondrial fraction (**Figure 3e**, Table S3, Tab 5), consistent with improvements in mitochondrial respiratory capacity typically observed following endurance training ^9, 47, 48^. While mitochondrial ROS-related proteins and pathways were acutely upregulated with exercise (**Figure 2e**), an overall downregulation of ROS-associated proteins was observed after 8 weeks of HIIT (**Figure 3f**, Table S3, Tab 6), consistent with a training-induced reduction in oxidative stress in skeletal muscle ^49^. Conversely, proteins related to mitochondrial dynamics ^50^, import machinery (e.g., TIMM/TOMM complexes) ^51^, chaperones and solute carriers ^52^ showed increased abundance, suggestive of training-induced mitochondrial network remodelling.

### Acute exercise mitochondrial proteostasis: Regulation of mitochondrial ribosomal, ROS, protein import, and mitochondrial dynamic proteins

Given the most robust proteomic remodelling observed was within the mitochondrial fraction (**Figure 2**), we further examined specific pathways and their respective proteins regulated by acute exercise, including those involved in translation, redox homeostasis, protein import and quality control systems. Cellular energetic stress can disrupt mitochondrial translation, respiration, protein import, or membrane potential, thereby activating conserved proteostatic mechanisms involving chaperones, proteases, protein import regulators, and mitochondrial dynamics ^53, 54^. Consistent with the observed downregulation of mitochondrial translation (GO:0032543) and mitochondrial gene expression (GO:0140053) by acute exercise (**Figure 2e**), the relative abundance of specific mitochondrial ribosomal proteins (mtRibo) decreased from PRE to POST, followed by recovery of protein abundance in this fraction after 3H (**Figure 4a**, Table S4). To determine whether this dynamic proteome profile was independent of overall mitochondrial content between timepoints, we applied an established mitochondrial protein enrichment (MPE) correction strategy ^7, 9^ (Table S6), which did not affect the outcome, indicating that the changes were unlikely driven by exercise-induced variations in mitochondrial fraction protein content (see **Supp. Figure 1c-e**). Visualisation of this coordinated clustering of mtRibo subunits showed both large (MRPL19, MRPL12, MRPL49, MRPL14) and small (MRPS25, MRPS35, MRPS36, MRPS11) subunits are downregulated from PRE to POST (Figure 4a). These time-course profiles (i.e., POST vs 3H) also showed mtRibo upregulation at 3H, suggesting commencement of restoration of mitochondrial translational capacity in the first three hours of recovery post-exercise (**Supp. Figure 1f**).

**Figure 4.**
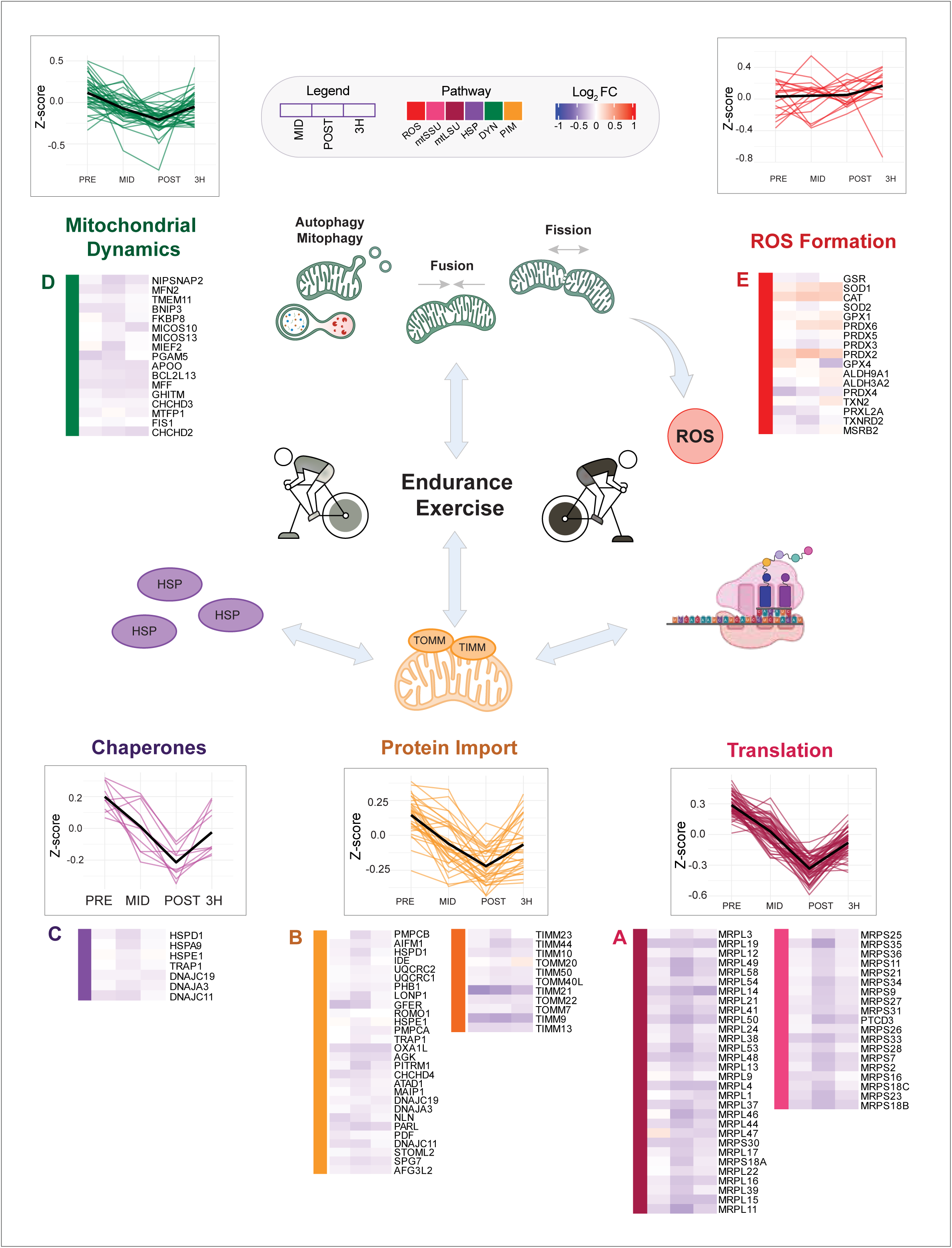
- Mitochondrial proteostasis pathways in response to acute exercise: regulation of mtRibo, ROS, protein import, and mitochondrial dynamics. Schematic of acute exercise-regulated mitochondrial pathways, including proteins involved with **a.)** mitochondrial translation/ribosomes (maroon), **b.)** protein import (orange), **c.)** chaperone proteins (light purple), **d.)** mitochondrial dynamics (dark green) and **e.)** reactive oxygen species (ROS) formation (red) as annotated by MitoCarta 3.0. Each pathway features scaled profile plots showing the relative abundance of each individual mitochondrial protein annotated to the specific pathway at each acute exercise timepoint (i.e., PRE, MID, POST and 3H), and the mean relative abundance (black) of the proteins within each pathway. Heatmaps of each differentially expressed protein within each specific pathway according to their adjusted P value (P <0.05) accompanies each profile plot, with the log2 fold change presented for each timepoint (MID, POST and 3H relative to PRE).

Protein import machinery (**Figure 4b**), chaperone proteins (**Figure 4c**) and proteins involved in mitochondrial dynamics (**Figure 4d**) exhibited similar temporal profiles. ROS formation related proteins generally increased during the time course of exercise (**Figure 4e**, Table S4). Proteins involved in mitochondrial protein import including TIMM21 and TIMM9 were progressively downregulated during/following acute exercise, while TOMM22, TOMM40L, and TOMM7 decreased at POST and partially recovered its mitochondrial abundance at 3H (**Figure 4b**). Mitochondrial chaperone proteins also exhibited coordinated regulation across the acute exercise time course (**Figure 4c**). Chaperones generally decreased from PRE to POST with partial recovery at 3H, consistent with transient perturbation of mitochondrial proteostasis during acute energetic stress. In parallel, changes in mitochondrial dynamics displayed a similar coordinated cellular stress-induced remodelling response (**Figure 4d**). MFN2, BNIP3 and FIS1 decreased POST and increased at 3H. MTFP1 showed a marked upregulation POST-exercise followed by a decline.

ROS-related proteins formed two main clusters that exhibited coordinated, but lower increases in magnitude over time. The first cluster included PRDX2, SOD1, PRDX6, and CAT, while the second comprised GPX1, ALDH9A1, and TXN2. GPX4 (glutathione peroxidase 4), which detoxifies lipid peroxidation and inhibits ferroptosis ^55^, rapidly increased at MID, followed by a rapid decrease post-exercise. In contrast, matrix-localised redox regulators such as SOD2 and GSR transiently decreased immediately POST-exercise but were increased after 3H (**Figure 4e**). Collectively, our findings illustrate coordinated temporal changes in mitochondrial protein regulation from mitochondrial translation, redox signalling, protein import, to dynamics following acute exercise, with evidence of increases in subcellular protein abundance beginning in the hours of post-exercise recovery, uncovering potential novel subcellular regulation of mitochondrial proteostasis underlying skeletal muscle adaptations to acute exercise.

### Exercise-induced ribosomal protein redistribution, stress adaptation, and translational regulation across muscle subcellular compartments

Given the robust mitochondrial protein remodelling observed post-exercise, particularly the decrease in mtRibo abundance (**Figure 4**), we further examined the subcellular responses of translation- and ribosome-associated proteins to acute exercise-induced energy stress. First, cytosolic translation-associated proteins exhibited phase-specific modulation in response to acute exercise. Initiation factors EIF1, EIF5A and EIF4E displayed increased protein abundance immediately POST, with a decrease at 3H. Elongation factors showed divergent responses: EEF1A2 and EEF2 abundance increased during/following exercise, whereas EEF1G and EEF1D decreased. Consistent with their known roles in contraction-mediated translational control, CAMK isoforms were predominantly increased after exercise ^56, 57^. The abundance of termination factors also dynamically changed in response to exercise, with GSPT1 decreasing at MID and increasing POST, and ETF1 increasing from MID to 3H (**Supp. Figure 3a**).

These cytosolic translational-associated protein signatures aligned with redistribution of ribosomal subunits across compartments (**Figure 5a**). In the nuclear fraction, ribosomal-associated proteins such as RPS18 and the ribosomal scaffold protein receptor for activated C kinase 1 (RACK1) increased post-exercise (**Figure 5b**) with RACK1 showing a modest but consistent increase (log□FC up to ∼0.22 at 3h post) and several ribosomal proteins were also detected in the mitochondrial fraction (**Supp. Figure 3b)**. In contrast, the abundance ribosomal proteins in the cytosolic fraction, including RPLP1 and RPSA, declined following exercise. RPS6KA3, a ribosomal S6 kinase subunit, increased at MID. In the mitochondrial fraction, small ribosomal subunits increased post-exercise, while large subunits increased at MID and subsequently declined post-exercise (**Figure 5a**). RACK1 was the most robustly increased protein following exercise in the mitochondrial fraction (**Figure 5b**), reaching a log□ fold change of up to ∼0.9 POST relative to PRE. We additionally assessed molecular chaperones (heat shock proteins, HSPs) in the nuclear fraction (**Supp. Figure 3c**). In the nuclear fraction, HSPs such as HSPB1, HSPB6, HSP90AA1 and HSP90B1 were increased following exercise. In contrast, other HSPs (e.g., HSPA1A, HSPA8, HSPA6, HSPA1L, HSPA2, HSPB7 and HSPB8) declined over time. Given the robust exercise-induced increase in RACK1 observed in the mitochondrial fraction, RACK1 proteomic data were next validated via targeted immunoblot analysis of the same fractionated lysates (**Figure 5c**). This confirmed the acute exercise-induced increase in RACK1 protein content in the mitochondrial fraction observed via MS, which was significantly increased immediately POST and at 3H post-exercise versus PRE (P < 0.05; **Figure 5d**), Trends for increased RACK1 protein in the nuclear fraction and decreased RACK1 protein in the cytosolic fraction were detected but failed to reach significance. (**Figure 5d**). Together these data demonstrate the utility of our subcellular proteomic dataset, confirming novel human muscle ribosomal regulation observed via MS, indicating a potential compartment-specific reorganisation of ribosomal and chaperone proteins in response to acute exercise.

**Figure 5.**
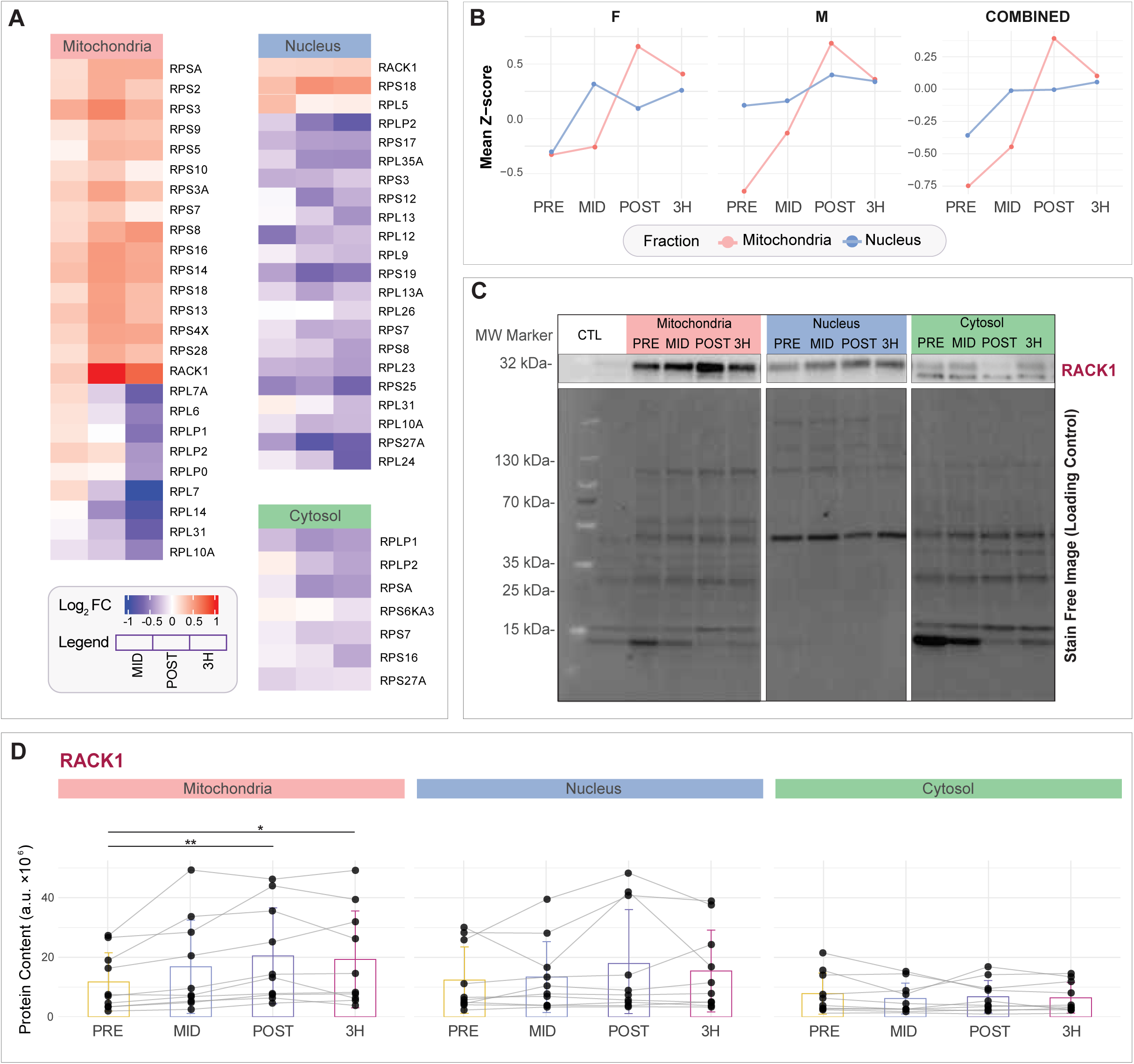
- Exercise-induced ribosomal redistribution, stress adaptation, and translational regulation across subcellular compartments. **a.)** Heatmap of differentially expressed ribosomal proteins (adjusted P < 0.05) identified in the mitochondrial, nuclear, and cytosolic fractions. Colours indicate log□fold-change at each acute timepoint (MID, POST, 3H relative to PRE). **b.)** Scaled profile plots showing mean z-score changes of Receptor for Activated C Kinase 1 (RACK1) detected in the mitochondrial and nuclear fractions across PRE, MID, POST, and 3H. Subpanels illustrate trajectories for females (F), males (M) and combined sexes (F), obtained from the Subcell-EX Shiny app for visualising compartment- and sex-stratified comparisons. **c.)** Representative immunoblot images of RACK1 (5 µg total protein loaded per lane, including molecular weight marker and pooled control sample in the first lanes) across subcellular fractions (mitochondria, nucleus, cytosol) at PRE, MID, POST, and 3H. Lane colouring reflects fraction identity: mitochondria (peach), nucleus (blue), cytosol (green). A Stain-Free total protein image is shown below as a loading control. **d)** RACK1 subcellular protein abundance in response to exercise, determined using band densitometry and expressed in arbitrary units (a.u.). Quantified data are presented as mean ± SD for a representative subset of n = 10 participants (5 males, 5 females). Comparisons are shown for PRE, MID, POST, and 3H within each fraction (mitochondrial, nuclear, cytosolic). (*P ≤ 0.05; **P ≤ 0.001) relative to PRE, determined by pairwise t-tests with Bonferroni correction.

Further to this targeted validation of RACK1, and to facilitate users’ further exploration of proteins identified/quantified in these subcellular proteomic datasets, we developed an interactive Shiny web application Subcell-EX (https://ereisman.shinyapps.io/subcell-ex/) that permits users to visualise exercise-induced changes in protein abundance across subcellular compartments and compare protein trajectories between sexes. Using this open access web tool, subcellular compartment-specific protein abundance changes can be visualised, such as the remodelling of RACK1 across the mitochondrial and nuclear fractions, with the more pronounced exercise-induced increases in RACK1 shown in the mitochondrial versus nuclear fraction, and in males versus females (**Figure 5b**).

## Discussion

Dynamic exercise induces significant cellular stress throughout contracting skeletal muscle tissues resulting in complex and coordinated responses across multiple subcellular compartments in response to both an acute bout of exercise and exercise training. Mitochondria, in particular, serve as key hubs for energy production, metabolic regulation, and coordinated responses to help cope with intracellular stress ^16^. Despite known roles of mitochondrial signalling and protein translocation between other organelles primarily established from whole muscle tissue analyses, major knowledge gaps remain in how exercise remodels the muscle proteome within and between specific subcellular compartments. This study provides the first insight into the subcellular proteomic remodelling that occurs in the mitochondria, nucleus, and cytosol following a single session of endurance exercise and training in human skeletal muscle from both males and females. Using subcellular fractionation combined with highly-sensitive MS technology, we identified distinct temporal and compartment-specific proteomic changes, with mitochondrial and nuclear fractions exhibiting more rapid and robust alterations than the cytosolic fraction. The relatively small cytosolic changes observed may reflect more transient and/or constrained responses to acute exercise, consistent with localised intracellular metabolic stress rather than broad proteomic restructuring across the cytosolic compartment ^58^.

Analysis of DEP revealed early regulation of transcriptional, translational, proteostatic, and metabolic pathways across compartments during and after exercise, indicative of a tightly coordinated, multi-compartmental response to help meet bioenergetic demands of exercise while maintaining proteome integrity. The mitochondrial proteome exhibited hallmark features of stress signalling, including signatures consistent with retrograde communication to the nucleus ^20, 59, 60^. These responses mirrored known mechanisms of proteome maintenance across subcellular compartments, involving coordinated regulation of mitochondrial chaperones, proteases, import machinery, and network remodelling pathways ^53^ ^54^.

Within three hours post-exercise, proteins involved in mitochondrial ribosomal assembly, import machinery, redox metabolism, and fission–fusion dynamics were significantly remodelled (**Figure 4**), suggesting that rapid proteomic reorganisation supports early mitochondrial network remodelling and bioenergetic adaptation. Downregulation of mitochondrial ribosomal components may reflect transient translational stress, whereas concurrent upregulation of redox-related proteins indicates exercise-induced oxidative signalling, a known trigger of transcriptional cascades that promote mitochondrial chaperones, import complexes (e.g., TIMM/TOMM), and dynamic remodelling ^61, 62, 63, 64, 65^. Consistent with this, we observed an initial decline in key components of mitochondrial translation, import machinery, and dynamics following acute exercise, reflecting a transient stress-induced suppression of protein translation in this fraction. These proteomic responses are characteristic of short-term adaptive intracellular regulation, in which non-essential mitochondrial biological processes are temporarily downregulated during acute metabolic stress and subsequently reinstated during recovery in the hours post-exercise, aligning with established models of stress-responsive proteostasis ^66, 67, 68^.

The patterns of ROS-related protein abundance during/following acute exercise also reflected their distinct sub-mitochondrial localisation: GSR and SOD2 localise to the matrix, while CAT and SOD1 associate with the intermembrane space, supporting the concept of compartmentalisation of redox signalling as a key regulatory mechanism that controls exercise metabolism in skeletal muscle ^69, 70^. These oxidative stress responses appear closely tied to mitochondrial quality control mechanisms, as ROS can impair protein folding, damage imported proteins and disrupt mitochondrial membrane integrity, all of which necessitate adaptive changes in import machinery and structural remodelling ^71^. Disruptions in mitochondrial protein import may therefore represent a key contributing factor underlying exercise-induced mitochondrial stress ^72^.

Of note was the marked downregulation of mtRibo with exercise that highlights a potential cellular stress-induced suppression of mitochondrial protein translation. This observed shift in ribosomal-associated protein dynamics aligns with a broader mechanism of mitochondrial stress adaptation, where the cell prioritises maintenance of other cellular energetic processes over protein synthesis ^73, 74^. The current findings in human muscle extend transcriptomic ^9, 75^ and proteomic studies ^76, 77^ by demonstrating that mitochondrial proteostasis is among the earliest detectable proteomic responses to exercise ^78, 79, 80^, and likely a critical step in initiating longer-term adaptations to endurance training. A rapid and selective regulation of key nodes of mitochondrial quality control was observed, including mtRibo proteins, TOMM/TIMM import machinery, redox enzymes and fission/fusion regulators. Concurrent changes in proteins involved in mitochondrial dynamics including OPA1, MFN1, and FIS1 further support this interpretation and align with prior observations linking mitochondrial network remodelling to exercise-induced cellular stress ^81^. Furthermore, we detected a relatively small, transient increase in the inner mitochondrial membrane protein MTFP1’s abundance in the mitochondrial fraction post-exercise, consistent with new signalling evidence of increased MTFP1 phosphorylation in response to an acute bout of HIIT ^10^ (**Figure 4**). These patterns suggest that mitochondrial impairment may be driven more by fission and fusion-related mechanisms than autophagy, with MTFP1 and other fission-related proteins potentially facilitating the removal of damaged mitochondria in response to oxidative stress. Recent ultrastructural studies also show that sprint interval exercise can induce mitochondrial disturbances before transcriptional responses emerge, supporting the idea that rapid mitochondrial dynamics indeed underpin early mitochondrial network remodelling ^81^.

Translational control is a highly regulated, energy-consuming process that underpins protein synthesis and cellular remodelling during stress and recovery. Recent evidence suggests that ribosomes may be specialised, with distinct regulatory roles depending on tissue type or cellular stress-inducing conditions ^82^. In the nuclear fraction, we identified a transient but coordinated enrichment of proteins involved in RNA processing, chromatin remodelling and transcriptional regulation within three hours post-exercise. These findings are consistent with transcriptional priming and extensive reports of increased transcription factor activity and stimulation of gene expression following acute exercise ^83, 84^. Most nuclear proteins returned to their baseline fractional protein abundance by POST, suggesting a tightly regulated, rapid increase in transcriptional regulation during exercise. In contrast, the cytosolic proteome exhibited fewer early changes, with only modest levels of proteome remodelling emerging POST, especially among proteins involved in folding, glycolysis and cytoskeletal organisation. This delayed cytosolic response may reflect downstream cytosolic programming following earlier mitochondrial and nuclear remodelling. Increased nuclear HSP protein abundance also aligns with stress adaptation pathways being triggered by misfolded proteins□with exercise ^85^.

An intriguing observation in the present study was the compartment-specific redistribution of ribosomal proteins in response to exercise. In the cytosolic fraction, a decline in ribosomal protein abundance (e.g., RPLP1 and RPSA) post-exercise may point to a temporary downregulation of bulk cytosolic translation. In contrast, small ribosomal subunits increased predominantly within the mitochondrial fraction, while RACK1 increased in both the mitochondrial and nuclear fractions. RACK1 is a multifunctional ribosome-associated scaffold protein involved in stress-responsive translational control, including ribosome quality control, sensing of ribosome stalling and selective initiation through interactions with EIF4E and EIF4A. Both of these proteins were also increased post-exercise in the cytosolic fraction ^86^ ^87^, and have been directly implicated in ribosome collision sensing and stalling-dependent quality control mechanisms ^88^. Notably, work in yeast has shown that the RACK1 homologue (CPC2) can anchor stalled ribosomes to the mitochondrial membrane, forming a hibernation-like, translationally inactive state during metabolic stress ^73, 89^, a concept further supported by recent evidence that RACK1 participates in stress-induced translational repression and ribosome pausing in mammalian cells ^90^. The post-exercise increase in RACK1 and small ribosomal subunits observed within the mitochondrial fraction may reflect a similar subcellular mechanism of translational pausing or ribosome tethering in human skeletal muscle, an adaptation that enables rapid reactivation of protein synthesis once cellular stress and energy homeostasis stabilises. These dynamic shifts in subcellular RACK1 protein abundance likely support skeletal muscle’s prioritisation of other muscle biological functions over translation to help maintain cellular resilience during exercise-induced energy stress ^91, 92^. To our knowledge, this is the first study to implicate RACK1 in human skeletal muscle adaptation to exercise, highlighting it as a candidate regulator of compartment-specific translational control from yeast to humans.

Considering increasing recognition of sex-based differences in several metabolic responses to exercise (i.e., patterns of substrate utilisation) ^93, 94, 95^, and the underrepresentation of females in human exercise studies ^96^, we examined whether sex influenced the physiological and subcellular proteomic responses to HIIT. While baseline sex-related differences in protein abundance remain biologically relevant, the largely absent sex-by-exercise effects observed across both acute and training phases suggest that subcellular proteomic adaptations are broadly conserved between sexes when matched for age and fitness. These findings indicate that mixed-sex cohorts can be informative for studying exercise-induced subcellular remodelling, while underscoring the importance of considering sex as a biological variable during study design and baseline characterisation.

By integrating differential centrifugation with time-resolved MS across multiple subcellular fractions, this study represents the most comprehensive subcellular proteomic analysis of human muscle molecular adaptations with exercise to date. Collectively, our datasets spanning acute exercise and training suggest that early subcellular proteomic responses may lay the molecular groundwork to initiate downstream adaptation in response to repeated exercise-induced stress. Extending beyond acute perturbations, training was associated with consolidation of subcellular proteomic remodelling, characterised by coordinated increases in proteins related to mitochondrial energy production, import machinery and organelle organisation. These changes are consistent with longer-term reorganisation of protein networks supporting enhanced oxidative capacity and proteostasis. The data highlight the mitochondrion’s central role in coordinating compartment-specific protein regulation of ribosomal subunits, translation factors, and cellular stress-associated proteins that collectively regulate proteostasis to maintain cellular homeostasis during and following exercise. As highlighted by the identification of RACK1 as a top exercise-regulated protein from the proteomic data and follow-up immunoblot validation, the temporal and spatial resolution of our human muscle subcellular proteomic datasets represents a valuable resource to identify novel exercise-regulated protein targets and guide future mechanistic studies aimed at decoding the molecular basis of exercise-induced adaptation. Finally, to facilitate widespread use of these datasets, we created an interactive user-friendly application enabling exploration of compartment-specific and sex-specific protein responses to exercise.

### Limitations of the study

Several limitations should be considered when interpreting the findings from this study. First, the use of subcellular fractionation enhanced resolution of proteomic responses to exercise by reducing overall muscle sample complexity, enabling the identification/quantification of less abundant proteins within mitochondrial, nuclear, and cytosolic fractions. However, fraction purity remains a consideration. While fractionation protocols typically utilise freshly collected tissue to preserve organelle integrity, our study applied an optimised protocol for frozen muscle samples, a practical necessity in large-scale human studies with prolonged recruitment and serial muscle biopsy sample collection windows. Snap-freezing and thawing muscle biopsies may compromise membrane integrity, potentially leading to leakage or redistribution of proteins across compartments, complicating interpretation of true subcellular localisation ^97^. Accordingly, while fraction enrichment was confirmed using established marker proteins (**Supp. Figure 1a–b**), proteins detected within a given fraction should be interpreted as compartment-enriched rather than exclusively localised. Cross-fractional protein identification remains an inherent challenge in subcellular proteomic analyses, particularly for proteins that can be localised in multiple intracellular compartments and/or undergoing dynamic intracellular trafficking. As such, our dataset is well suited to examining compartment-specific changes in protein abundance in response to exercise, rather than inferring precise intracellular localisation or protein translocation.

Second, the participant cohort was limited to healthy, young, previously untrained females and males. As such, the generalisability of these findings to other populations, such as older adults, individuals with chronic disease, or highly trained and athletic populations remains unclear. Additionally, menstrual cycle phase was not standardised in female participants, however, current evidence suggests this influence is limited in the context of acute exercise responses ^98, 99, 100^. In addition, the absence of a non-exercise control group represents a limitation, as PRE samples served as baseline. While inclusion of such a control group would increase participant and logistical burden, it may help further disentangle exercise-induced changes from biopsy-related effects such as local inflammation in future studies ^101, 102^.

Finally, while subcellular fractionation enabled total proteomic analysis of compartment-specific adaptations, this sample preparation workflow required substantial human muscle biopsy starting material and therefore limited our ability to assess multi-omics layers such as phosphoproteomics. Our proteomic analysis represents a snapshot of muscle subcellular protein abundance at a given timepoint and does not directly assess post-translational modifications, protein turnover, or dynamic protein translocation. Integration of complementary approaches, including phosphopeptide enrichment or spatial proteomic methods, represents an important direction for future work. Despite these limitations, the clear temporal and spatial distinctions observed across subcellular compartments provide strong evidence for a tightly coordinated, compartment-specific proteomic remodelling in response to acute exercise, highlighting the value of subcellular proteomics for identifying exercise-responsive proteins and pathways relevant to human skeletal muscle plasticity, underlying exercise’s wide health benefits.

## Author contributions

D.J.B., J.A.H., N.J.H conceptualised the study. E.G.R., D.F.T., D.J.B., J.A.H., and N.J.H. devised the study methodology. E.G.R. and D.F.T. delivered the exercise sessions and performed sample collection. E.G.R. performed subcellular fractionation and MS sample preparation. Proteomic analysis was performed by E.G.R. and D.Y. with input from C.H., N.J.C. and N.J.H. E.G.R. and N.J.C. performed statistical and bioinformatic analysis and visualisation. E.G.R., J.A.H, and N.J.H. wrote the initial manuscript draft, and all authors revised the manuscript. All persons designated as authors qualify for authorship, and all those qualifying for authorship are listed. All authors have read and approved the final version of the manuscript.

## Acknowledgements

This research was funded by an Australian Research Council (ARC) Discovery Project grant (DP200103542) awarded to D.J.B., J.A.H. and N.J.H. This research was also supported by an Australian Government Research Training Program (RTP) Scholarship https://doi.org/10.82133/C42F-K220 awarded to E.G.R. and MS infrastructure funding from the Victorian Higher Education State Investment Fund (VHESIF; awarded to Australian Catholic University and St Vincent’s Institute).

## Resource availability

All source data used to interpret, verify, and expand upon this research are included with this paper. The raw mass spectrometry proteomics data have been deposited to the ProteomeXchange Consortium through the PRIDE partner repository with the dataset identifier PXD063881. (Reviewer login details - username; password: UOf60TedTdaC).

## Lead contact

Correspondence to Dr Nolan J. Hoffman

## Methods

### Experimental design and subject details

#### Ethics approval

This study was approved by the Victoria University (HRE20-212) and Australian Catholic University (2022-2518R) Human Research Ethics Committees, registered with the Australian New Zealand Clinical Trials Registry (ACTRN12621001062819), and conformed to the standards set by the 1964 Declaration of Helsinki (and its later amendments and comparable ethical standards). All participants received detailed study information, outlining the potential benefits and risks of all study procedures before providing written informed consent.

#### Study participants

Forty healthy, untrained participants, including 20 males and 20 females (26.5□±□5.9 years; 173□±□10□cm; 70.8□±□13□kg; BMI 24.9□±□3.4; V□O_2_max 31.1 ± 5.6 mL/min/kg; mean□±□SD), volunteered and completed the acute exercise phase of the study, with all participants included in the final analysis. Sample size (n) was determined using scikit-learn machine learning algorithms ^103^ and prior clinical proteomics studies ^104^, indicating that 14 participants per sex provided sufficient power (≥80%) to detect significant effects of exercise (P□<□0.05) and model subcellular proteomic network adaptations potentially linked to mitochondrial biogenesis.

#### Study design and exercise testing

Participants were instructed to refrain from intense physical activity for at least 48 h before each performance assessment (96 h for muscle biopsy procedures) and abstain from consuming caffeine for a minimum of 3 h prior. All tests were conducted at a similar time of day to limit potential circadian rhythm-related variability.

##### Graded Exercise Test (GXT)

Participants performed a GXT before and after the 8-week training program to assess lactate threshold (LT) and associated power output (□LT), defined as the point where blood lactate concentration increased exponentially during exertion. The test consisted of 4-min stages on an electronically braked cycle ergometer (Lode Excalibur Sport, Groningen, The Netherlands), with resistance increasing by 7.6% per stage until cadence dropped below 60 RPM. Initial intensity was 20% of estimated peak power (W peak), based on demographic, anthropometric and activity data as previously described ^105^. Blood samples were collected at rest and after each stage via antecubital vein cannula (∼ 1 mL), or capillary sampling when required. Lactate was analysed using an automated analyser (YSI 2000 Glucose/Lactate Analyser, YSI, Ohio, USA), and LT was determined using the modified Dmax method ^105^.

Gas exchange was measured every 15-s using a MOXUS Metabolic Cart (AEI Technologies, Pennsylvania, USA). V□O□peak was calculated from the highest two 15-s averages. Heart rate and rating of perceived exertion (RPE, Borg RPE 20 scale ^106^) were recorded during the final 30 s of each stage.

##### Ramped Exercise Test (RXT)

Before training, participants completed an RXT to assess maximal power output (□max), which may be underestimated by the GXT. The test consisted of 1-min stages for 8–12 min total, starting at 20% of estimated □max and increasing power output until cadence dropped below 60 RPM or volitional fatigue. Protocols were customised to elicit fatigue at ∼10 min based on participant demographics, body composition, and activity levels ^105^. After a 5-min rest, a verification bout at 105% of W□max was performed to confirm V□O□max and □max. As in the GXT, expired gases were measured every 15 seconds, and V□O□max was determined from the two highest consecutive readings. Heart rate (Polar Electro Oy, Kempele, Finland) and RPE (Borg RPE 20 scale^106^) were recorded in the final 10 s of each stage.

#### Physical activity and nutritional controls

Participants recorded their dietary intake for three days, including the 24 h preceding each experimental trial, and were instructed to replicate the same diet before each GXT/RXT. To minimise variability in whole-body and muscle energy availability, all participants received a standardised meal during the biopsy trial day and for the 12 h following the initial HIIT session. Energy intake was individualised using the Mifflin–St Jeor equation ^107^ with an activity factor of 1.2, with additional energy provided to account for exercise energy expenditure (calculated from time × work). Standardised meals were aligned with the *Australian Dietary Guidelines* and provided at a macronutrient composition of approximately 50% carbohydrate, 30% fat, and 20% protein.

#### Muscle biopsies and exercise protocols

A total of five muscle biopsies were obtained from each participant throughout the study under local anaesthesia (5 mg.mL^-1^ Lidocaine) from the *vastus lateralis* using the suction-modified Bergström muscle biopsy technique. After baseline testing and refraining from intense physical activity for at least 72 h, a resting biopsy (PRE) was collected from the vastus lateralis in the overnight fasted state and immediately snap-frozen. Participants then performed a single HIIT session (4 × 4-min intervals, prescribed as training intensity = W□LT + 0.45(W□max -W□LT), with two-min rest periods). Additional biopsies were collected after the 2nd interval – MID, immediately post-exercise (POST) and 3 h after the PRE biopsy. All participants were then assigned to eight weeks of HIIT, which required them to perform the same HIIT session described above 4 days/week for 8 weeks total with progressive increases (∼2.5%/week) in the training load (final training intensity = W□LT + 0.75(W□max -W□LT)). A fifth biopsy was collected at least 96 h following the final training session. To minimise the influence of local inflammatory responses, each muscle biopsy was obtained through a separate incision spaced at least 1 cm from the previous site. To control for potential effects of repeated biopsies, the first four (acute) muscle biopsies were collected from the same leg, while the final biopsy was obtained from the leg selected by the participant. All samples were snap-frozen to preserve protein integrity.

#### Muscle subcellular fractionation

A ∼50 mg portion of each skeletal muscle biopsy collected during the first HIIT session (PRE, MID, POST, 3H) and post-training were subjected to subcellular fractionation using stepwise differential centrifugation into mitochondrial, nuclear, and cytosolic fractions. Each biopsy sample was homogenised in 1.3 mL SEMH buffer (220 mM mannitol, 70 mM sucrose, 1 mM EDTA, 20 mM HEPES-KOH, 1:100 phosphatase/protease inhibitor cocktail (PIC), Cell Signaling Technology) using a 2 mL glass dounce homogeniser (60 strokes on ice; Sigma-Aldrich, St. Louis, Missouri, USA). This protocol was adapted from prior skeletal muscle methods to improve protein detection and throughput for MS ^108, 109^.

The homogenate was centrifuged at 200 *g* (4□°C) to remove intact cells and debris. If the pellet exceeded 50% of tube volume, an extra 30 strokes were performed on ice. The resulting supernatant was centrifuged at 1000 *g* for 10 min (4□°C) to separate the unpurified mitochondrial/cytosolic supernatant and unpurified nuclear pellet.

The unpurified mitochondrial and cytosolic supernatant was centrifuged at 2000 *g* for 10 min (4 °C), and the resulting supernatant was spun at 12,000 *g* for 15 min (4 °C) to isolate the purified mitochondrial fraction pellet, which was then resuspended in SDS lysis buffer (comprised of 50 mM triethylammonium bicarbonate (TEAB), 5% SDS (w/v)). The cytosolic fraction was isolated by centrifuging the supernatant at 21,100 *g* for 15 min at 4 °C.

To obtain the nuclear fraction, the unpurified nuclear pellet was resuspended in 500 μL SEMH buffer and homogenised by pipetting up and down gently before centrifugation at 1000 *g* for 10 min (4 °C). The resulting pellet was resuspended in 300 μL NET buffer (20 mM HEPES-KOH, 0.5 M NaCl, 20% glycerol, 1.5 mM MgCl□, 0.2 mM EDTA, 1% Triton X-100, 1:100 PIC) and homogenised by pipetting, followed by tip probe sonication at 30% amplitude (3 × 10 s bursts, 10 s rest). Benzonase (0.5 µL) was added to the isolated nuclear fraction, to further facilitate release of all nuclear proteins and enhance protein yield ^110^. The resuspended nuclear pellet was spun at 21,100 *g* for 15 min (4 °C), resulting in the final nuclear fraction supernatant. Proteins from the nuclear and cytosolic fractions were precipitated in acetone (4 x sample volume) at – 20□°C overnight, then pelleted (21,100 *g*, 1 h, 4□°C), and resuspended in SDS lysis buffer for subsequent analyses.

Protein quantification of each isolated fraction was performed using Pierce™ bicinchoninic acid (BCA) protein assay (Thermo Fisher, Waltham, Massachusetts, USA).

#### Immunoblotting – Subcellular fractions

To assess the purity of subcellular fractions isolated from frozen muscle biopsies, a subset of samples from each isolated fraction was randomly selected and analysed by immunoblotting. Protein pellets were solubilised in SDS lysis buffer denatured for 10 min at 95 °C and mixed with 4 x Laemmli buffer solution (pH 6.8, 250mM Tris-HCl, 8% SDS, 20% glycerol, 0.03% bromophenol blue, 10% β2-mercaptoethanol). Lysates (5 μg protein per lane) were loaded onto Bio-Rad Stain-Free™ Criterion™ TGX (4%–20%) gels (Bio-Rad, Hercules, California, USA) and proteins were separated using SDS polyacrylamide gel electrophoresis (SDS-PAGE).

Proteins were transferred to polyvinylidene difluoride (PVDF) membranes using the Trans-Blot® Turbo™ semi-dry system (Bio-Rad, Hercules, California, USA), then blocked with 5% bovine serum albumin (BSA) and 0.1% TWEEN-20 detergent in tris-buffered saline (TBS; TBS-T)). Membranes were washed in TBS-T and incubated overnight with rolling at 4°C with primary antibodies of each subcellular compartment protein marker diluted in TBS-T. Antibodies used to verify compartment enrichment/purity included anti-histone (H3; Abcam #Ab1791; nuclear marker, 1:3000 dilution in 5% BSA), anti-mitochondrial heat shock protein 70 (mtHSP70; Invitrogen #MA3-028; mitochondrial marker, 1:1000), and anti-lactate dehydrogenase (LDHA; Cell Signaling Technology #2012; cytosolic marker, 1:1000). Receptor for Activated C Kinase 1 (RACK1, Cell Signalling Technology, #5432 CST) was also interrogated via immunoblotting to validate MS results. Following at least 3 x 10 min washes in TBS-T with vigorous rocking, membranes were incubated with HRP-conjugated secondary antibodies (goat anti-mouse IgM, Thermo Fisher #31172; goat anti-rabbit IgG, CST #707), diluted 1:5000 in 5% BSA. Chemiluminescent signal was detected using Clarity™ Western ECL (Bio-Rad Hercules, California, USA) or SuperSignal™ West Femto (Thermo Fisher), and blots were imaged on a ChemiDoc™ MP system (Bio-Rad). Protein visualisation and densitometry were performed using Image Lab software (version 6.1, Bio-Rad). The volume density of each protein band at the target molecular weight was quantified and first normalised to an identical pooled lysate sample loaded onto each gel, then further normalised to the total protein detected per lane using stain-free imaging technology (**Supp. Figure 1**).

#### Proteome sample preparation

Derived from methods previously outlined by Zougman *et al.* ^111^, fractionated samples underwent the following preparation steps. Samples were lysed in 1 x SDS lysis buffer (comprised of 50 mM triethylammonium bicarbonate (TEAB) and 5% SDS (w/v)) at 95°C for 10 min, then sonicated for 20 min. Disulfide bonds were reduced with 10 mM tris-2-carboxyethylphosphine (TCEP) and samples were again heated at 95°C for 10 min. Alkylation was performed by adding chloroacetamide (CAA, Sigma-Aldrich) to a final concentration of 40 mM and incubating at room temperature for 30 min.

SDS lysates were acidified by adding 12% phosphoric acid (H□PO□) at a 1:10 ratio, yielding ∼1.2% final concentration, crucial for protein filtration at this pH. Next, 350 µL of S-Trap buffer (pH 7.1; 90% Methanol, 100 mM TEAB, C_7_H_17_NO_3_) was added to the acidified lysate to induce colloidal protein formation. The mixture was loaded into a 96-well S-Trap™ plate (Protifi, Fairport, New York, USA) and centrifuged at 1500□*g* for 2□min or until all solution passed through, trapping the proteins within the column. Flow-through was collected in a 2 mL deep 96-well plate. Captured proteins were washed three times with 200□µL S-Trap buffer and centrifuged for 2 min at 1500 *g* or until all solution passed through. Proteins were digested overnight (∼16□h) at 37□°C with trypsin/LysC (1:25 ratio) in 125□µL of 50□mM TEAB (i.e., digestion buffer). Peptides were sequentially eluted using 80 µL of digestion buffer, then 80 µL 0.2% aqueous formic acid (FA) for each elution, with subsequent centrifugation at 4000 *g* for 1 min. Hydrophobic peptides were recovered with a final elution of 80 µL 60% (v/v) acetonitrile (ACN; C□H□N) containing 0.2% FA, with all elutions from the same sample pooled. Peptides were lyophilised in a vacuum concentrator and re-constituted in 0.1% TFA and 2% ACN. Samples were then clarified by centrifugation at 21,300 x *g* prior to MS analysis.

#### Liquid chromatography-MS/MS data acquisition

Peptides underwent reverse-phase high-pressure liquid chromatography (HPLC) at acidic pH and were separated using a Dionex UltiMate™ 3000 RSLCnano. Peptides were then injected onto a 2 cm PepMap™ trap cartridge (Thermo Fisher) and separated using μPAC™ Neo HPLC columns (180 μm × 500 mm, Thermo Fisher). The mobile phases were composed of buffer B 80% ACN containing (0.1% formic acid; FA) and buffer A (0.1% FA). A gradient was applied at 40 °C to the mobile phase over 50 min; starting with buffer B at 2.5%, increasing to 5% at a flow rate of 700 nL/min over 5 min. Then it increased to 35% at a reduced flow rate of 350 nL/min until 35 min, followed by a sharp increase to 45% until 38 min. Subsequently, the gradient increased to 95% at 41 min, where the flow changed back to 700 nL/min. Finally, it returned to 2.5% until 46 min before completing the run at 50 min.

An Orbitrap Lumos Fusion MS instrument (Thermo Fisher) equipped with a NanoSpray Flex ion source (Positive Ion (V): 2000, Ion Transfer Tube temperature 275 °C) was utilised under the following conditions: MS1 scan was acquired within the range of 300–1250 m/z (at a resolution of 120,000, with a target 4e5 AGC and 40% RF Lens, and an injection time of 50 ms) followed by MS2 in DIA mode with higher energy collisional dissociation and with orbitrap detection (OT HCD). MS2 scan range was 145-1450 m/z, 15,000 OT resolution, activation HCD, 33% HCD collision energy, 22 ms maximum injection time, 14 m/z quadrupole isolation width, normalised AGC Target 1000%, and absolute AGC Value 5e5. The acquired raw DIA data were analysed using the integrated software suite DIA-NN (v 2.0), with processed data exported to csv file format and subsequently analysed in RStudio (v2024.09.0+375).

#### Bioinformatic analysis of proteomics data

##### Clean up and normalisation

Proteins identified as contaminants were removed, with only high confidence proteins (with more than one unique peptide) included. Proteins were further filtered to remove those that contained >30% missing values across participants. Remaining missing values were imputed using the k-nearest neighbour (knn) method from the *impute* package (v1.78.0) in R (v4.4.1; k = 10). Data were then normalised via variance stabilisation normalisation (vsn) using the *vsn* package (v3.72.0).

##### Differential expression analysis

Differential expression analysis was conducted using the *limma* package (v.60.6) ^112, 113^ with empirical Bayes moderation via the eBayes function. To account for the paired design (i.e., repeated measures on the same individuals), blocking was performed using the duplicateCorrelation function. Proteins with an adjusted P-value < 0.05 (Benjamini–Hochberg method) were considered differentially expressed. Differentially expressed proteins were visualised in heatmaps using unsupervised hierarchical clustering (Euclidean distance, ‘average’ linkage) via the *ComplexHeatmap* package (v.2.20.0) in R (v4.4.1). Enrichment analysis was performed using the *enrichR* (v3.2) ^114^ and *clusterProfiler* (v.4.12.6) packages, with terms considered enriched at adjusted *p* < 0.05 (BH method). Mitochondrial proteins were annotated using the MitoCarta 3.0 database ^32^. Data visualisations were generated using *ggplot2* and associated R packages. All code developed for this manuscript is available on GitHub: https://github.com/reisman-eg/MitoX-SubCell.

### Statistical Analysis

Statistical analysis of physiological and blood parameters was performed using the rstatix package (v.0.7.2). Paired Student’s t-tests were used for within-subject comparisons before and after the 8-week training intervention (PRE vs POST), with significance set at P < 0.05. To assess the effects of sex and time, two-way repeated-measures ANOVA was performed. Post-hoc pairwise comparisons were conducted only for relevant contrasts (e.g., PRE vs POST within each sex), without adjusting for unrelated comparisons, to identify specific group differences. Additionally, pairwise t-tests (Bonferroni-adjusted) were used to analyse immunoblots to compare mean protein abundance relative to PRE within each subcellular fraction (mitochondrial, nuclear, and cytosolic).

### Subcell-EX interactive web application

To enhance user accessibility and engagement with the subcellular proteomic dataset, we developed Subcell-EX, an interactive R Shiny web application (https://shiny.rstudio.com). The application enables users to explore temporal protein profiles across mitochondrial, nuclear, and cytosolic fractions, with the option to visualise one or two proteins simultaneously. Users can toggle between combined or sex-specific views and examine differential expression for each subcellular compartment. The application also includes a simplified gene ontology (GO) enrichment module, allowing users to input custom gene lists and assess fraction-specific biological processes. All code for Subcell-EX is available at https://github.com/reisman-eg/MitoX-SubCell.

## Supplementary Figure Legends

**Supplementary Figure 1.**
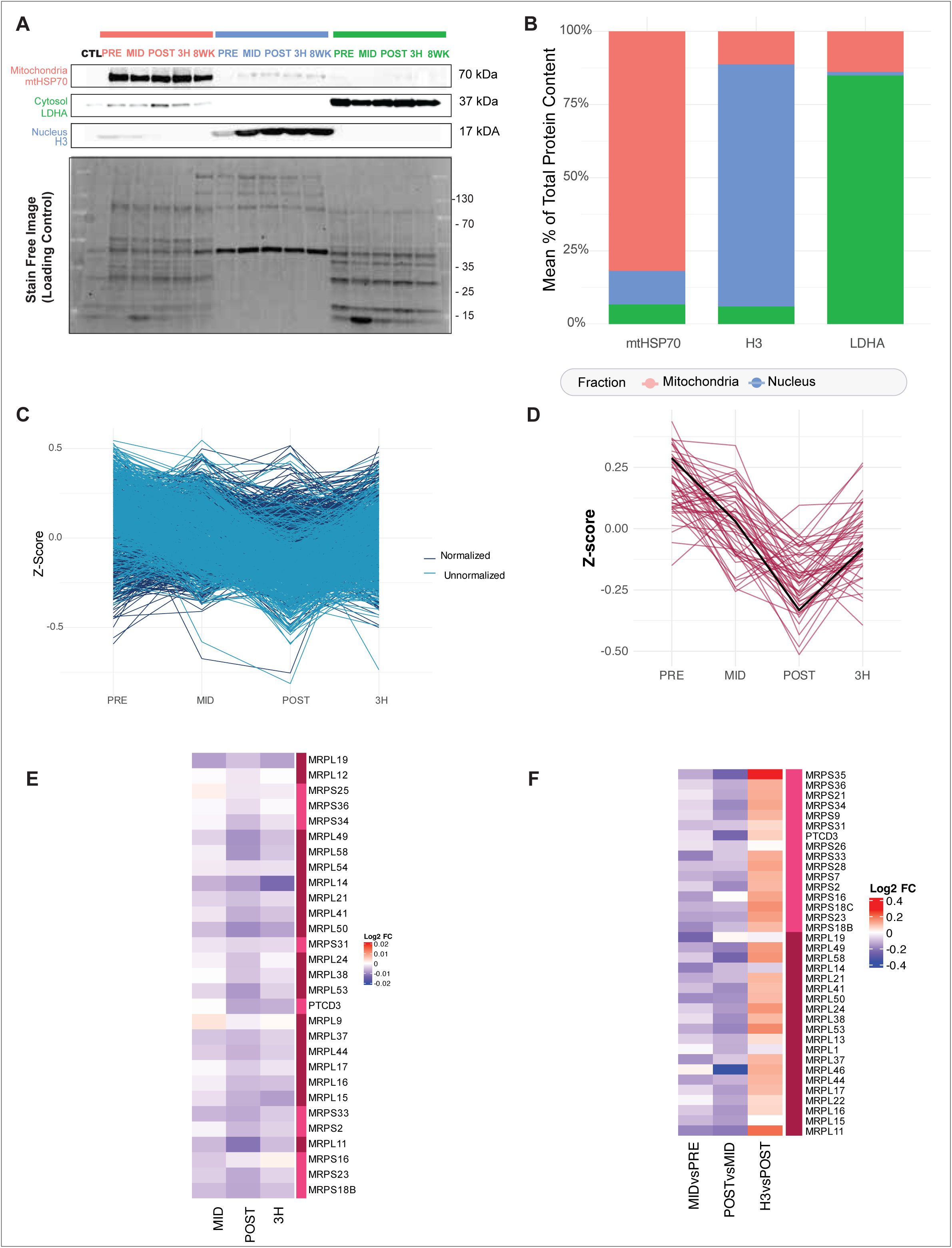
**a.)** Representative immunoblot images demonstrating benchmarking of subcellular fraction purity using antibodies specifically targeting a single subcellular compartment marker (mtHSP70; mitochondria, LDHA; cytosol, H3; nucleus). Lanes: CTL – pooled control sample; Mitochondria (Peach): PRE, MID, POST, 3H, 8WK; Nucleus (Blue): PRE, MID, POST, 3H, 8WK; Cytosol (green) PRE, MID, POST, 3H, 8WK. Stain Free Image (Loading Control) presented below. **b.)** Purity analysis of the yield of subcellular marker proteins. The percentage shown is obtained by the ratio between western blot volume density values for mitochondrial heatshock protein 70 (mtHSP70) n = 10, Histone (H3) n =10, and lactate dehydrogenase (LDHA) n =8 in an individual target compartment divided by the sum of the volume density values present in all compartments in each subcellular fraction. A representative subset of individuals were selected for PRE, MID, POST, 3H & 8WK. **c.)** Scaled profile plot demonstrating data corrected for mitochondrial protein expression versus uncorrected data **d.)** Scaled profile plot demonstrating mtRibo protein data corrected for mitochondrial protein expression **e.)** Heatmap displaying differentially expressed proteins according to the adjusted P value of mtRibo proteins corrected for mitochondrial protein expression presented with log_2_ fold change for each timepoint relative to PRE. **f.)** Heatmap displaying differentially expressed proteins according to the adjusted P value of mtRibo proteins presented with log_2_ fold Change as a time course for MIDvsPRE, POSTvsMID, 3HvsPOST.

**Supplementary Figure 2.**
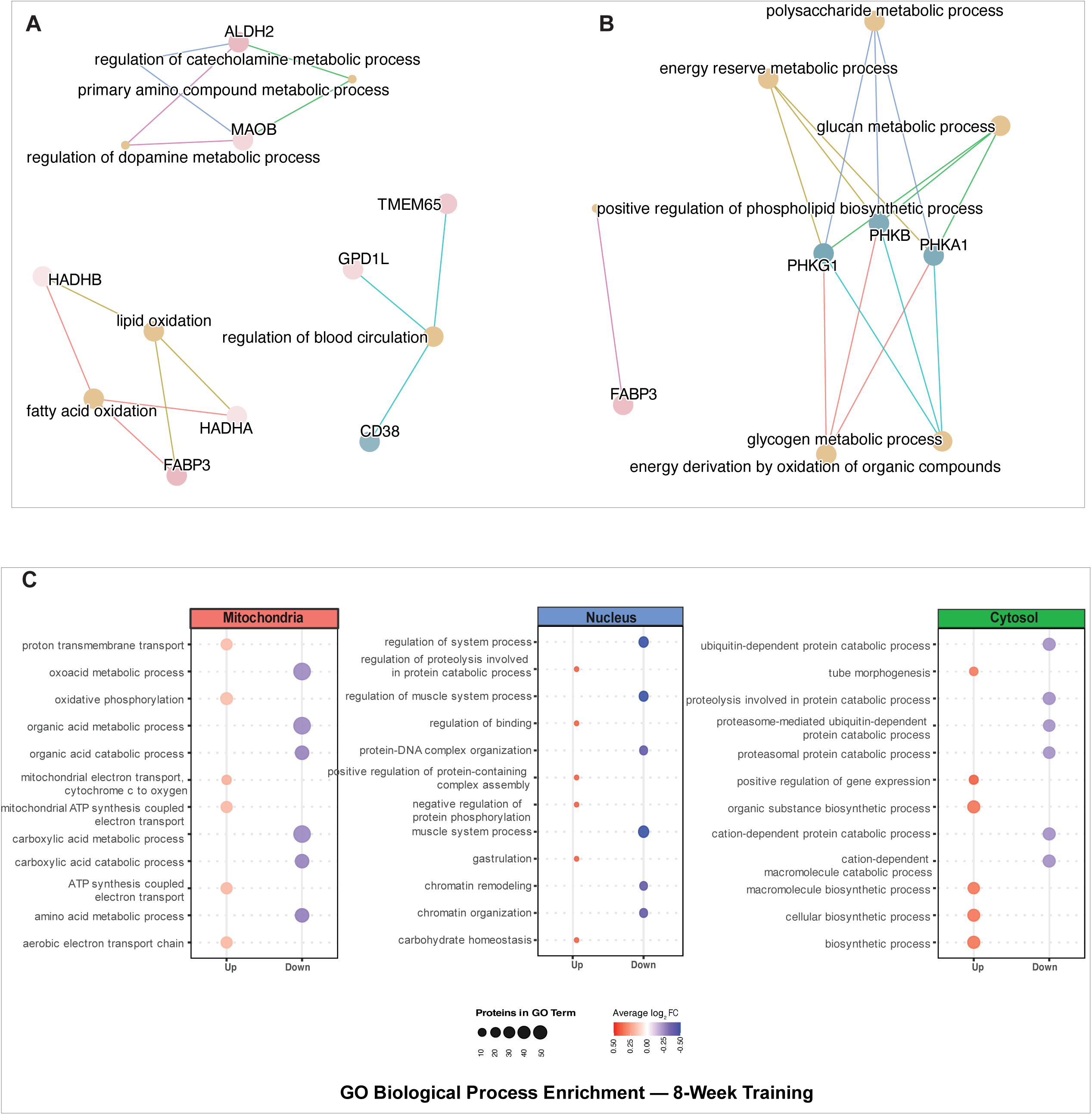
**a.)** *ClusterProfiler* network plot using differentially expressed (Benjamini-Hochberg adjusted P value < 0.05) terms for sex differences identified across each timepoint in the mitochondrial fraction as per Supp. Figure 2a. Pathway terms were identified using Gene Ontology Biological Processes (GOBP). Colours indicate in which sex (pink – females, blue – males) proteins identified from a specific GOBP term were higher. **b.)** *ClusterProfiler* network plot using differentially expressed (Benjamini-Hochberg adjusted P value < 0.05) terms for sex differences identified across each timepoint in the cytosolic fraction. Pathway terms were identified using GOBP. Colours indicate in which sex (pink – females, blue – males) proteins identified from a specific GOBP term were higher. **c.)** EnrichPlot dot plot analysis of differentially expressed proteins following 8 weeks of training, annotated as mitochondrial (Mitocarta 3.0 ^32^) or as nuclear/cytosolic (Human Protein Atlas ^36^). Pathway enrichment was performed using GOBP. For each timepoint, the top eight enriched terms (ranked by adjusted P-value) are shown for upregulated and downregulated proteins. Dot size represents the number of proteins associated with each term, while colour saturation reflects the average log□ fold change for that timepoint and direction of regulation.

**Supplementary Figure 3.**
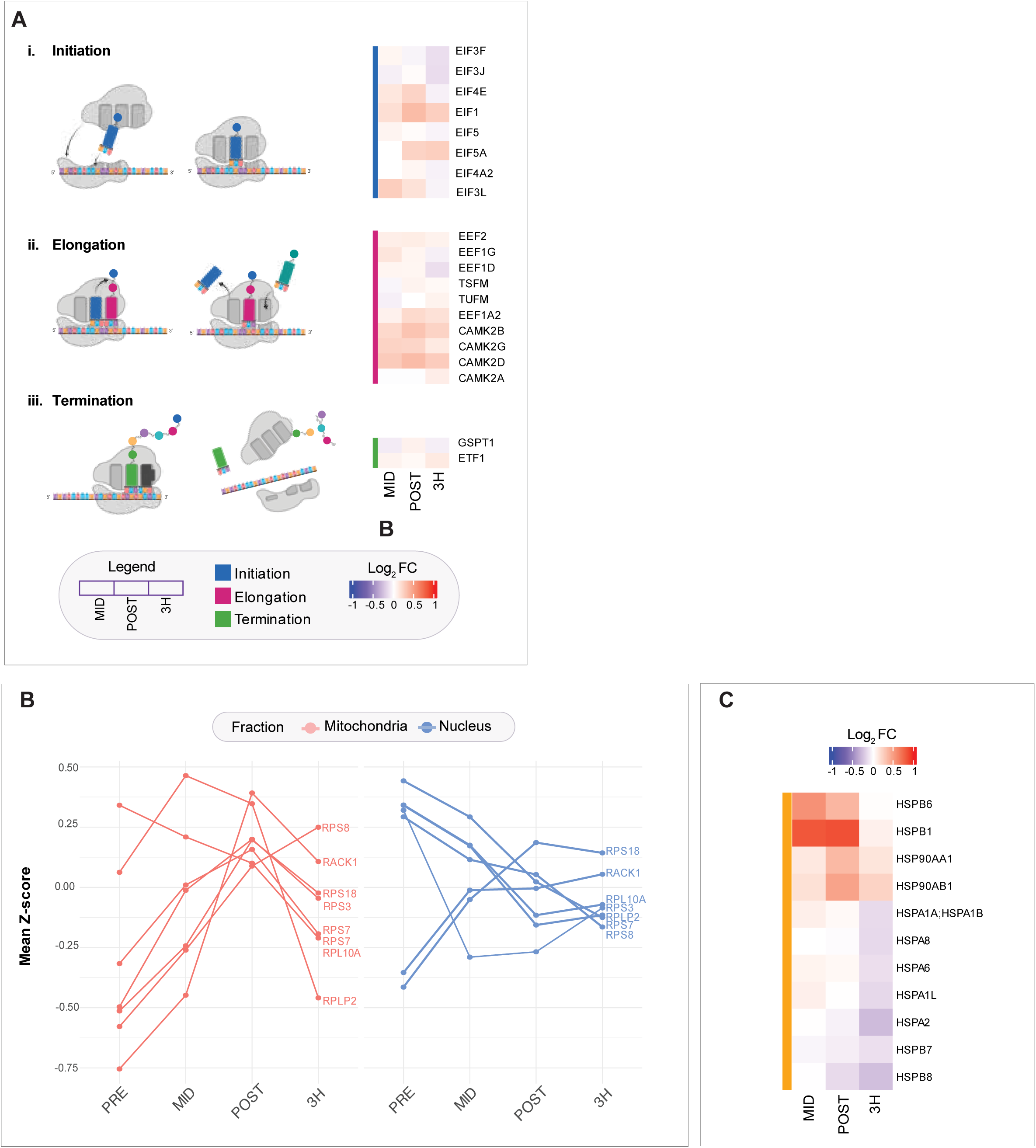
**a.)** Heatmap displaying differentially expressed proteins according to the adjusted P value of proteins associated with phases of translation including **a)** initiation, **b)** elongation and **c)** termination alongside a schematic of protein translation (created with https://BioRender.com). **b.)** Scaled profile plot demonstrating the relative responses of ribosomal proteins identified in both the mitochondrial and nuclear fraction **c.)** Heatmap displaying differentially expressed proteins according to the adjusted P value of heat shock proteins presented with log_2_ fold change for each timepoint relative to pre in the nuclear fraction.

## Description of Additional Supplementary Files

### File name: Supplementary Table 1

Description: Participant characteristics, Raw intensities for mitochondrial, nuclear and cytosolic proteomic data, and log intensities for all identified proteins in all samples following the normalisation described in the methods for mitochondrial, nuclear and cytosolic proteomic data. Data also for immunoblot targeted analysis of representative samples of mitochondrial, nuclear and cytosolic fractions using markers mtHSP70, H3 and LDHA

### File name: Supplementary Table 2

Description: The normalised Data (Log2) with *limma* calculations for mitochondrial, nuclear and cytosolic proteomic data of MID vs PRE, POST vs PRE, 3H vs PRE & 8WK vs PRE. GO Biological Processes (GOBP) results for mitochondrial, nuclear and cytosolic proteomic data for proteins annotated as mitochondrial (according to MitoCarta 3.0) in the mitochondrial fraction, nuclear (according to Human Protein Atlas) in the nuclear fraction and cytosolic (according to Human Protein Atlas) in the cytosolic fraction using enrichplot & clusterProfiler. Top 6 GOBP for up or down regulated proteins at each time point is presented

### File name: Supplementary Table 3

Description: Physiological data for VO2max, lactate threshold (Dmod, WLT) and peak power output (Wpeak) is presented along with measures of mitochondrial content (CS activity and MitoVD). The normalised Data (Log2) with *limma* calculations for mitochondrial, nuclear and cytosolic proteomic data of 8WK vs PRE. Scaled intensities for known mitochondrial proteins for all samples annotated as complexes CI-CV by MitoCarta from PRE to 8WK post training in the mitochondrial fraction. Protein functional classes presented with the scaled means (z-scores) for each functional group (defined by MitoCarta 3.0) from PRE to 8WK post training in the mitochondrial fraction.

### File name: Supplementary Table 4

Description: Scaled intensities for known mitochondrial proteins for all samples at acute time points, PRE, MID, POST, 3H annotated as functional group (defined by MitoCarta 3.0) protein import (PIM), mitochondrial dynamics (DYN), chaperones (CHA), reactive oxygen species (ROS) in the mitochondrial fraction. Scaled z-scores for mitochondrial ribosomal proteins (mtRibo) for PRE, MID, POST, 3H as defined by Mitocarta 3.0 in the mitochondrial fraction are also presented.

### File name: Supplementary Table 5

Description: Description: The normalised Data (Log2) with *limma* calculations for mitochondrial, nuclear and cytosolic proteomic data of MID vs PRE, POST vs PRE, 3H vs PRE & 8WK vs PRE including sex as a factor for males vs females (MvsF) in response to exercise. Within timepoint MvsF is also calculated.

### File name: Supplementary Table 6

Description: Mitochondrial protein expression (MPE) calculation is subsequently presented for identified mitochondrial proteins within the mitochondrial fraction. Data is then normalised and corrected for mitochondrial content for identified mitochondrial proteins within this fraction. Subsequent normalised corrected Data (Log2) with *limma* calculations for mitochondrial proteomic data of MID vs PRE, POST vs PRE, 3H vs PRE of the acute phase. Further analysis of uncorrected data (Log2) with *limma* calculations for mitochondrial fraction calculated as a time-course i.e. MID vs PRE, POST vs MID, 3H vs POST is presented.

